# Unsupervised Graph-Based Learning Predicts Mutations That Alter Protein Dynamics

**DOI:** 10.1101/847426

**Authors:** Robert L. Peach, Dominik Saman, Sophia N. Yaliraki, David R. Klug, Liming Ying, Keith R. Willison, Mauricio Barahona

## Abstract

Proteins exhibit complex dynamics across a vast range of time and length scales, from the atomistic to the conformational. Adenylate kinase (ADK) showcases the biological relevance of such inherently coupled dynamics across scales: single mutations can affect large-scale protein motions and enzymatic activity. Here we present a combined computational and experimental study of multiscale structure and dynamics in proteins, using ADK as our system of choice. We show how a computationally efficient method for unsupervised graph partitioning can be applied to atomistic graphs derived from protein structures to reveal intrinsic, biochemically relevant substructures at all scales, without re-parameterisation or *a priori* coarse-graining. We subsequently perform full alanine and arginine *in silico* mutagenesis scans of the protein, and score all mutations according to the disruption they induce on the large-scale organisation. We use our calculations to guide Förster Resonance Energy Transfer (FRET) experiments on ADK, and show that mutating residue D152 to alanine or residue V164 to arginine induce a large dynamical shift of the protein structure towards a closed state, in accordance with our predictions. Our computations also predict a graded effect of different mutations at the D152 site as a result of increased coherence between the core and binding domains, an effect confirmed quantitatively through a high correlation (*R*^2^ = 0.93) with the FRET ratio between closed and open populations measured on six mutants.

## 1 Introduction

Proteins undergo dynamics at different scales that allows them to inter-convert between conformations [1]. Such motions are effectively encoded in their molecular structure: fast and local vibrations develop collectively into the motion of secondary structures and, eventually, larger conformational domains [1]. Despite the biological importance of such dynamics in mediating function (including catalysis, allostery and mechanical processes), there is a lack of concrete evidence supporting conclusively different theories that attempt to link structure, dynamics and function. In this paper, we present the application of a data-driven, unsupervised, graph-theoretic methodology[2] to analyse atomistic structural graphs of proteins in order to identify parts of a protein that display coherent behaviour at different time scales. We then compare our computational predictions to experimental FRET measurements of domain motion obtained from wild-type and mutated ADK.

Domain motions usually occur on relatively slow time scales, with kinetic barriers between states on the order of several *kT*. The rates of inter-conversion are usually monitored using either single-molecule FRET [3] or hydrogen-deuterium exchange combined with either mass spectrometry or Nuclear Magnetic Resonance (NMR) [4]. In some situations, Laue X-ray diffraction [5] has also been informative.

Molecular dynamics (MD) simulations using atomistic force fields are a powerful technique to study protein dynamics [6, 7]. However, the long time scales of catalytic motions (microseconds to milliseconds) are currently out of computational reach for larger proteins. In general, characterising the effect that small perturbations have on long-term dynamics of proteins require long simulations, which remain computationally expensive despite recent advances [8]. Furthermore, the diversity of force fields used in MD has prevented a consensus on some of the underpinning physical drivers of protein dynamics.

In parallel to dynamical simulations, a variety of discrete approaches for the analysis of protein structures have been developed [9, 10, 11, 12]. Normal mode analysis (NMA) is often used to characterise the molecular fluctuations of elastic network models (including gaussian network models and anisotropic network models) and to identify domain motions in a number of systems [9]. However, NMA is limited as a result of the harmonic approximation and is often constrained to backbone atoms which inherently ignore important inter-subdomain interactions [10]. Another network method to explore dynamics of proteins is the Structural Perturbation Method (SPM) which explores the propagation of phonons for intra-protein signalling and has provided insights into the allosteric mechanism in GroEL [11].

In previous work, we have introduced efficient graph-theoretic methodologies that retain key physico-chemical details through an energy-weighted atomistic protein graph, whilst effectively probing dynamical substructures of proteins in a number of biological scenarios [2, 13, 14, 15]. Here, we extend this graph-theoretical approach to perform systematic rankings of full mutational scans in an unbiased manner, scoring the mutations according to their effect in disrupting the multi-scale domain structure of the protein on time scales accessible to FRET measurements. The computational analysis of alanine and arginine *in silico* scans allows us to select candidate mutations, which we then test experimentally using FRET. This cycle of computational predictions and experimental validation is applied to Aquifex ADK, a protein that balances molecular stability and large domain motions in the pursuit of cellular energy homeostasis [16]. Our results demonstrate a strong quantitative correlation between changes in population of FRET states and computational mutational predictions.

## Results

### Unsupervised identification of biologically relevant graph partitions at multiple scales

The computational analysis of the structure of ADK (closed conformation, PDB ID:2RGX) using Markov Stability (MS) [2, 17, 18, 19, 13, 14, 20] is presented in Fig. 1. From protein data bank files (PDB) [21], we construct a fully atomistic graph of the protein to ensure that the collective effect of all interactions is considered (see Methods and Fig. S1). Markov Stability then explores all levels of spatial and temporal resolution to identify robust graph communities encoded in the atomic protein structure in an unsupervised manner. To achieve this, MS exploits the time evolution of a diffusive process on the protein graph to identify groups of atoms that behave similarly over a particular time scale in response to perturbation inputs. We identify relevant partitions as being both highly reproducible (i.e., signalled by dips in the average Variation of Information (VI) [22] of the ensemble of solutions) and temporally persistent (i.e., signalled by long plateaux in time) [13, 19] (see Methods).

**Figure 1:**
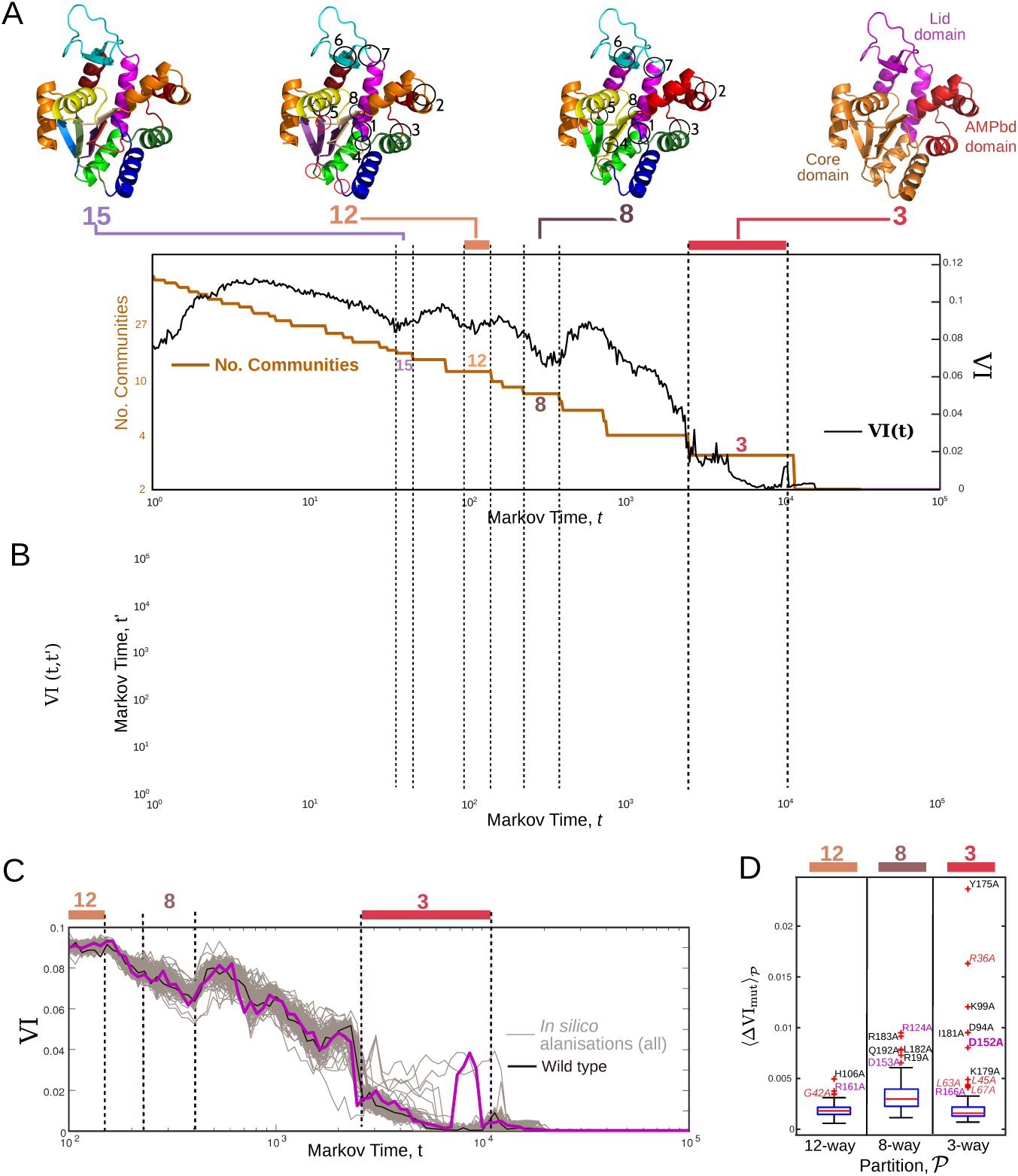
(a) Markov Stability (MS) results for the atomistic graph of ADK (closed conformation, PDB ID: 2RGX). MS finds optimised partitions at different levels of resolution as scanned by the Markov time, *t*. The similarity between partitions is measured using the variation of information 0 ≤ VI ≤ 1, where VI = 0 indicates equality of partitions. We seek partitions that are both persistent over time (i.e., optimal over a long plateau of *t*) and robust under the optimisation of the cost function (i.e., they exhibit a dip in VI(*t*)). At long time scales we find four robust large-scale partitions into 15,12, 8 and 3 communities, which are shown mapped onto the protein structure. The 15-way partition is highly coincident with secondary structures. The 12-way and 8-way partitions match the domains and hinges highlighted by Kern and coworkers [1] (black circles), with additional boundaries (circled in red) not previously identified which may also act as hinges. The highly robust 3-way partition agrees with the catalytic subdomains (lid, core, binding) reported in the literature. Fig. S2 presents the MS scan extending to the short scales together with the scan of the open conformation. (b) A heatmap of VI(*t, t′*) where the optimal partition at each *t* is compared with the optimal partitions at all other values of *t*. The diagonal of the heatmap, VI(*t*_*i*_, *t*_*i*_), will have a value of 0 where we compare the same partition to itself. Large blue blocks correspond to similar partitions over a long time period (temporally robust) and therefore may have some biologically meaningful structure. (c) MS results for all *in silico* alanisations of the 202 amino acids of ADK, one at a time (grey lines) together with the wild type (black line). Most mutations have a negligible effect on the robustness of the different partitions but some of the mutations induce changes, especially in the 3-way partition. (d) Box and whisker plot of the average change in the robustness of the 12-, 8- and 3-way partitions induced by all 202 alanine mutants. The red line in each box indicates the median, and the box indicates 25/75 percentiles. The outliers (plotted with red crosses) are the mutations that induce large changes at each timescale and coloured according to their domain: lid (purple), AMP_bd_ (red, italic), core (black). The mutant D152A is the only high scoring mutation affecting the 3-way partition in the lid subdomain.

The robust partitions found by MS correspond to biologically relevant levels of organisation across a wide range of scales: from chemical groups, amino acids and biochemical motifs (i.e., the common building blocks of all proteins) to the larger subdomains that are characteristic to each protein [13, 14, 23]. The multi-resolution analysis of two ADK structures (open and closed) in Fig. S2 show that the small scale partitions are highly coincident across both structures, whereas changes in their robustness appear at the longer time scales beyond secondary structures.

Figure 1A focuses on the long time scales associated with ADK conformational dynamics, starting with the 15-way partition into communities that correspond to secondary structures (alpha helices and beta sheets). The hierarchy of robust partitions of increasing coarseness shows 12-way and 8-way partitions that correspond closely to communities separated by hinges defined by Henzler-Wildman and coworkers [24], yet our method finds four additional community boundaries that might be flexible hinges previously not identified in ADK (red circles Fig. 1A). A similar set of community boundaries has been shown to be important in identifying epitopes in allergenic proteins [23], thus cementing the dynamical relevance of the subgraphs identified by MS.

At long time scales, MS finds a robust, persistent 3-way partition which agrees with the catalytic subdomains defined in the literature (Fig. 1A). Specifically, the three communities found (30-71; 112-170; rest of the residues) match well the canonical subdomains reported in the literature: AMP_bd_ (31-69), lid (114-170), core (remaining residues).

### *In silico* alanisation identifies mutants that disrupt partitions on time scales observable by FRET

The numerical efficiency of the MS methodology allows us to carry out a full *in silico* alanine scan of the ADK structure: each individual residue of the protein is substituted by alanine, one at a time, and the full MS analysis is recomputed for each mutant. Mutations were introduced using PyRosetta [25] and energy minimisation was performed using a short (200 ps) MD GROMACS simulation [26]. Once the mutation is introduced and a modified atomistic graph is obtained, the multi-resolution organisation of the mutant is recomputed with Markov Stability (see Methods).

As expected, most alanine mutations induce very small changes in the observed partitions (and their robustness), yet a few mutants elicit large variations. In order to facilitate experimental comparisons, we quantify the effect of mutations on the large-scale partitions associated with dynamical processes observable via FRET. Fig. 1B shows that the robustness of the 12-way and 8-way partitions are relatively unaffected by mutations, whereas a few mutations (e.g., D152A) reduce the robustness of the 3-way partition drastically, as seen by the large loss of robustness (i.e., increase in VI) they induce. All mutants are ranked according to their effect on each partition 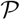 by computing the loss of robustness 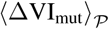 against the wild type (see Methods) over the corresponding time scale, as shown by box plots in Fig. 1C.

The outliers in Fig. 1C correspond to uncharacteristically disruptive mutations at different time scales of biological relevance. We identify H106A, R161A and G42A (for the 12-way) plus R124A, R183A, L183A, Q192A, R19A and D153A (for the 8-way) as high ranking mutants that disrupt the domain structure related to hinges. Of those, H106A, R183A, L182A, Q192A and R19A lie in the core subdomain away from the active site, whereas G42A lies in the AMP_bd_ subdomain. D153A, R124A and R161A are located in the lid subdomain, and together they form a hydrogen bond network.

Markov Stability identifies a series of high ranking mutations that affect the 3-way partition into lid/core/binding subdomains thought to move independently during the catalytic process [27] and which can be probed with FRET. Of these residues, only D152 is in the lid subdomain, and, in accordance with the dynamic nature of the lid [24], it becomes a focus of our investigation.

D152 is found on the 3/10 *α*-helix 7 at the hinge between the floppy lid and lid-connector helices. Although it is highly conserved across ADK species, D152 has not been previously linked to the dynamics of substrate binding. On closer inspection, D152 forms a salt bridge with the high scoring R36 in the AMP_bd_ domain, suggesting that this interaction is important for ADK conformational dynamics. Within the lid subdomain, D152 forms hydrogen bonds with R161 (high scoring in the 12-way partition) and R150, which has been identified experimentally as playing a dual role in the structural stabilisation and catalytic activity of *E. Coli* ADK [28]. Furthermore, D152 is an immediate neighbour of D153, which is similarly conserved and identified as disruptive for the hinges of the 8-way partition. All considered, these observations lead us to select D152 as a candidate for our experimental mutational FRET studies, as a previously unidentified residue outside the binding region, which induces strong effects at the large conformational scales probed with FRET whilst being coupled to the shorter scales associated with hinge flexibility.

### Single-molecule FRET confirms the dynamical changes induced by D152 mutations

To test the relevance of our predictions to the dynamics of the lid, we carried out single-molecule FRET (smFRET) experiments on D152 mutants of ADK. The distance between the lid and AMP_bd_ domains is monitored using the FRET between a donor (Alexa 488) and acceptor (Alexa 633) located on the respective domains (Fig. 2A). (For details of FRET pair preparation and measurements, see Methods.) The smFRET efficiency (*E*) was calculated from the total number of donor and acceptor photons counted during the dwell time of the laser focal volume. The apparent FRET efficiency was measured in the absence of substrate, so as to focus on population shifts solely induced by mutations. The FRET efficiency histograms were fitted to a two-state distribution mixture model to determine the relative proportions of the open and closed conformations (Fig. 2B and Methods).

**Figure 2:**
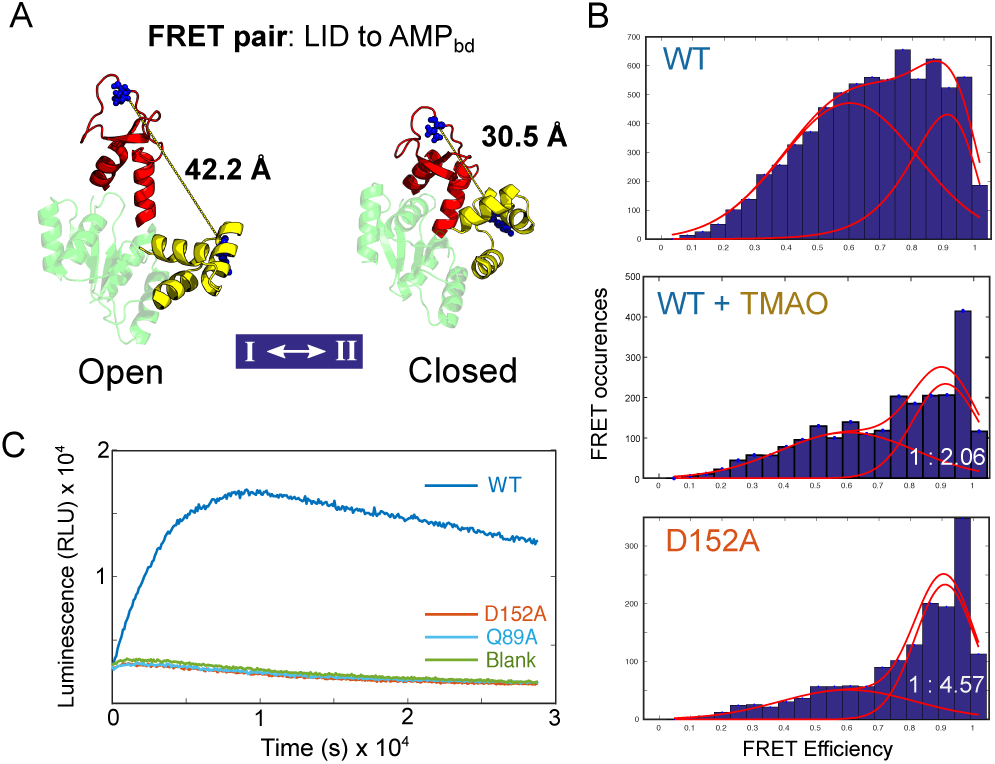
(a) FRET pair constructed to probe the dynamics between lid and AMP_bd_ domains in ADK [24]. (b) Relative FRET efficiencies of: wild type, wild type in the presence of osmolyte (TMAO, 3M), and D152A mutant. (c) A luciferase activity assay shows the inhibition of D152A relative to wild type, blank and a control mutant (Q89A, dead catalytic site).

The FRET efficiency histogram of the wild type exhibited an open (*E* = 0.56) and a closed (*E* = 0.82) state sampled to a similar fraction (1: 0.91). When the osmolyte TMAO was used to induce a closed conformation, the population shifted towards the closed state in a proportion (1: 2.06), in agreement with FRET measurements by Henzler-Wildman *et al* [24]. Likewise, the mutant D152A also exhibited a large shift in FRET efficiency towards the closed state in proportion (1: 4.57), suggesting that this mutation significantly alters the free energy of the closed conformation. Enzyme activity assays revealed complete catalytic inhibition for the D152A mutant, despite D152 not being a binding residue (Fig. 2C). These results suggest that Asp152 plays a critical role in the structural integrity of the lid domain and is essential for catalysis despite not binding substrate, highlighting the long-range, slow-time effects that single mutations can induce.

### Computational analysis of different D152 mutants predicts experimental FRET population shifts

To exploit and validate the atomistic capabilities of the MS computational methodology, we explored the effect of other D152 mutations on the robustness of the partition of interest and on FRET measurements. Our calculations show that the mutations have different effects on the robustness of the 3-way partition (see Fig. S3 for the MS calculations of the full set D152 mutants). Fig. 3A shows the MS computations for six D152 mutants, chosen to reflect a range of effect sizes: alanine (A) induces the largest loss of robustness (i.e., a large increase in VI(*t*) over the relevant time scale), whereas glutamic acid (E) has minimal effect. Figure 3B shows the smFRET experiments for the same six D152 mutants, measured and fitted as above. As in our computations, the six mutations exhibit a large variation in the FRET populations: from an almost equal open:closed ratio for D152E (1:1.24) to a large shift towards the closed conformation in D152A (1:4.57). Figure 3C shows that the closed to open population ratio is strongly correlated (*R*^2^ = 0.93) with the loss of robustness of the 3-way partition, i.e., increase of 〈VI〉_3-way_ in our MS computational analysis. Intuitively, this reflects the biophysical notion that the opening and closing motion of the enzyme is reliant on the presence of three *well-defined* blocks articulated by hinges. When the modularity of the structure is diluted, the closed conformation becomes prevalent, possibly through local unfolding or collapse of the protein structure, as suggested by the loss of catalytic activity for all D152 mutants, including D152E (Fig. S4). Overall, these results highlight the importance of Asp152 for structure, dynamics and catalytic function.

**Figure 3:**
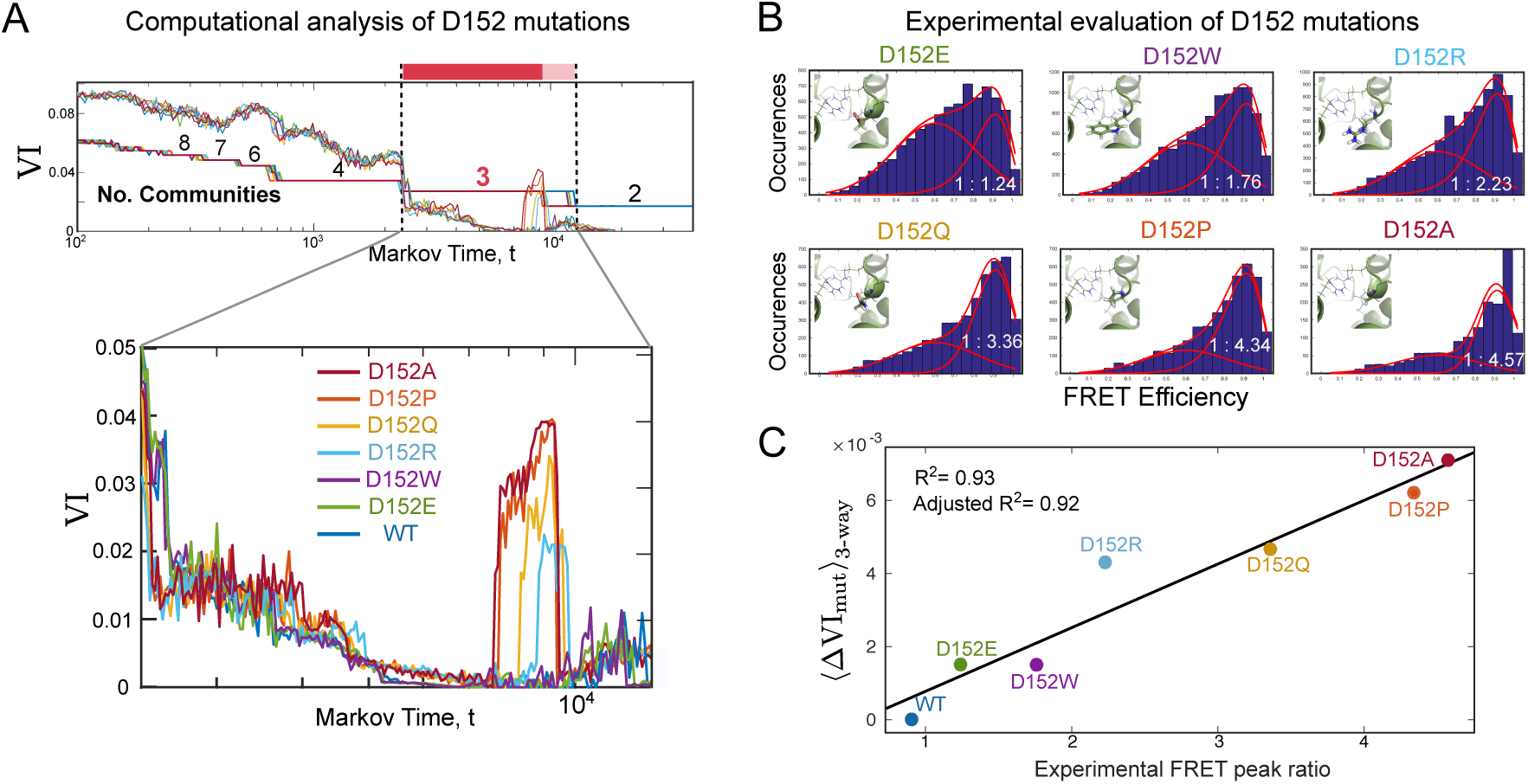
(a) Markov Stability results for six D152 *in silico* mutants, showing increasing loss of robustness of the 3-way partition. For the full set of D152 mutations see Fig. S3 in the SI. (b) As a result of each mutation, the FRET histograms exhibit a graded shift in the population ratio between open and closed states. The FRET histograms are fitted with a normal and log-normal distribution as described in Methods. Insets show a local view of the chemical structure of each D152 mutation. (c) The experimental population shift towards the closed state in smFRET is correlated with the loss of robustness of the 3-way partition 〈∆VI_mut_〉_3-way_ computed with MS, which reflects the loss of independence of the core and AMP_bd_ domains.

To understand further the loss of robustness of the 3-way partition in D152 mutants, we examined in more detail the computational outcomes of the atomistic MS analysis. As described in Methods, the VI(*t*) measures the average similarity of the partitions found through 1000 optimisations of the Markov Stability cost function at each Markov time. If the optimisation finds always the same partition, then VI = 0 and the partition is robust. If the algorithm finds very different solutions, then VI is large reflecting the existence of different, yet viable, structural partitions that coexist in the ensemble of solutions. Since VI is a proper metric [22], the ensemble of partitions can be represented in terms of a geometric landscape: the histogram of frequencies can be plotted as a function of the 2D projection of the distance between partitions obtained with multidimensional scaling (MDS). Figure 4 presents this projected landscape of partitions for the wild type, D152R and D152A—three structures with increasing value of 〈VI〉_3-way_. We extract 1000 partitions from the wildtype, D152R and D152A each at the Markov time where we observe significant differences in VI as shown in Figure 3A. The MDS allows us to visualise how similar the total set of 3000 partitions are to each other. The wild type overwhelmingly (99.6%) exhibits a partition (P_1_) that corresponds to the three standard subdomains (core, AMP_bd_, and lid). For D152R and D152A, partition P_1_ becomes less prevalent, appearing only in 82% and 35% of the optimisations, respectively, as these mutant structures have access to another locally optimal partition (P_2_), which mixes parts of the core and AMP_bd_domains. The alternative partition P_2_ appears 17% of the time for D152R and, in increasingly dissimilar form, 65% overall for D152A (Fig. 4). Hence the loss of robustness of the 3-way partition is the consequence of beta sheet *β*-3 and alpha helix *α*-5 in the core becoming grouped with the AMP_bd_ domain, with hinges 1, 4 and 8 losing their role as connectors between both domains. Our results suggest that the shift of the FRET population towards a closed state is linked to the core and AMP_bd_ subdomains becoming more fused and less modular.

**Figure 4:**
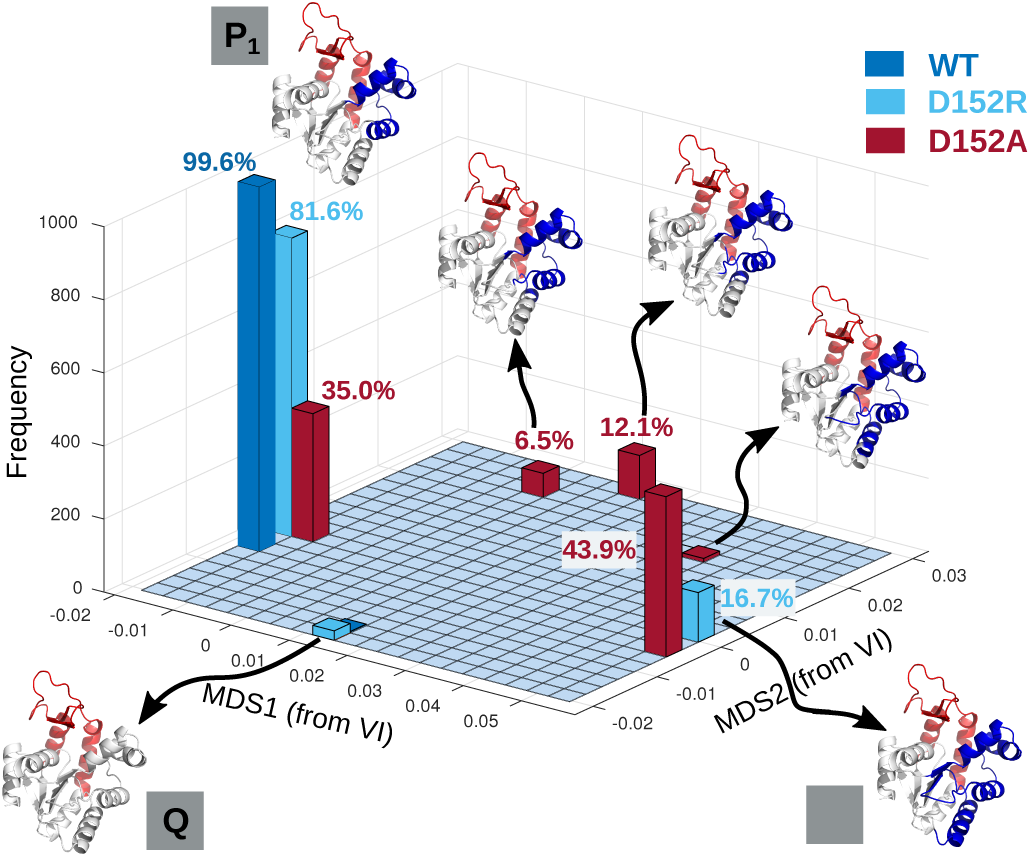
The loss of robustness of the 3-way partition by substitution at the D152 site is related to a reduction in the independence between the core and AMP_bd_ domains, as shown by partitions found by MS when analysing the wild type, D152R and D152A mutants. We use multidimensional scaling (MDS) to embed in two dimensions (MDS1 and MDS2) the distance VI, which is a proper metric, between the partitions found by 1000 MS optimisations. The P_1_ partition corresponds to the three standard subdomains (lid, AMP_bd_, core); the P_2_ partition has a large area of the AMP_bd_ domain joining the core. The 2-way partition *Q* has AMP_bd_ and core fused. The wild type is overwhelmingly found in the well-partitioned, highly modular P_1_; D152R partially accesses the partially fused P_2_; and D152A is found predominantly in P_2_ and on the transitional pathway between P_1_ and P_2_.

### Computational arginine scan identifies distinct mutations and functional effects

The results above exemplify how the MS computational framework not only detects the effect of local mutations on the large scale organisation of the protein, but can also capture the subtleties of chemical substitutions. Arginine residues are known to play an important role in the orchestration of enzymatic activity in ADK [29], and the importance of arginines in stabilising folds has long been purported [30, 31]. Our *in silico* alanisation identifies several significant arginine (R) residues in ADK (Fig. 1C). For example, R124, R161 and R166A, which form hydrogen bonds with D152 and D153, score highly relative to other lid mutations.

To test the effect of mutations that add, rather than eliminate, chemical bonds, we carried out a full computational arginine scan of ADK: mutating each residue, one at a time, to arginine; followed by MD re-equilibration of the structure; and a full MS analysis of the structure of each mutant, as described above. Figure 5A shows the box plot of the effect of all R mutations on the 3-way partition. In contrast to alanisation (Fig. 1C), the outliers in the arginine scan identify more residues in the lid and AMP_bd_ as significant hint at the importance of hydrophobicity and polarity in maintaining the large scale and core structure of the protein. The arginine scan also identifies D152R as a high scoring mutation, further highlighting the importance of the Asp152 residue. Other high scoring lid mutations are V157R, K160R and V164R. V164 is located on the lid connector helix, and has a number of interactions with the AMP_bd_ domain, specifically a strong hydrogen bond with Glu57 (Fig. 5B, top). This mutation has thereby the potential to restrict (or ‘staple’) the opening and closing motion associated with catalytic activity in ADK—hence we choose this mutant as a target for our experimental studies.

**Figure 5:**
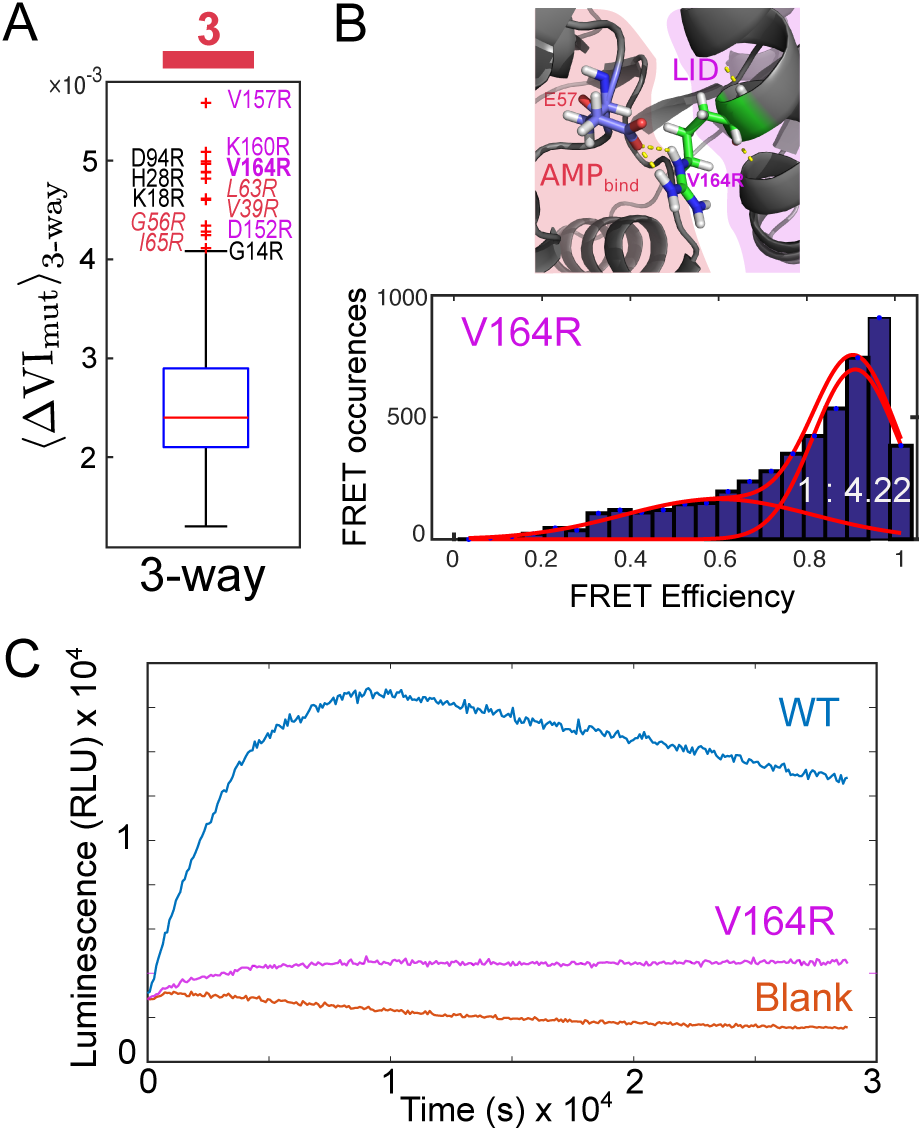
(a) A full *in silico* arginine scan of ADK identifies V164R as a significant outlier in disrupting of the 3-way partition. Box and whisker plot as described in Fig. 1C. (b) The smFRET of the V164R mutant exhibits a large population shift towards the closed state. Mutation of V164R introduces two strong hydrogen bonds between the lid and residue E57 in the AMP_bd_ domain. (c) V164R exhibits significant reduction, but not full inhibition, of the catalytic activity.

We tested experimentally this idea by constructing, labeling and carrying out smFRET measurements on the V164R mutant, as described in Methods. The smFRET measurements (Fig. 5B) showed a significant population shift towards the closed state, in accordance with the correlation in Fig. 3C. Importantly, however, this shift towards a closed conformation in V164R has distinct structural and functional effects. Structurally, the effect of the V164R mutation is to shift alpha helix 4 from AMP_bd_ to the core domain (Fig. S5). Functionally, whereas all D152 mutants (Fig. S4) are inactive, V164R exhibits catalytic activity, albeit at a significantly decreased rate (Fig. 5C). This reduction in activity is consistent with the slowing of the opening and closing of the catalytic site due to the formation of bonds between lid and AMP_bd_, similar to the effect observed by Kovermann *et al* [32] by adding a disulphide bond.

The population shift in the closed-to-open ratio (from 0.91 in WT to 4.22 in V164R) correlates with the 3.5-4 fold reduction in the catalytic rate (Fig. 5C). However, there is no justification that changes in the equilibrium constant can necessarily lead to an equally measureable effect on catalysis. Whilst our results agree with Kovermann *et al* [32] who found that an arrested closed conformation reduced catalytic activity, we can’t fully conclude that there is a causal relationship. Aviram *et al* [33] have shown that there is a relationship between the domain opening and closing times that can measurably modulate enzymatic activity, however, this doesn’t extend to the open and closed equilibrium.

## Discussion

We have shown how multi-scale partitioning of atomistic energy-weighted graphs obtained from protein structures can reveal, in an unsupervised manner, relevant scales of structural organisation, from chemical groups through secondary structures and hinges to large scale conformational groupings. The method can be used to score systematically the effect of mutations on protein organisation at different length and time scales. We applied our analysis to the enzyme ADK, carrying out full computational alanine and arginine scans and found mutations that disrupt strongly the large scale 3-way partition linked to the lid/binding/core domains.

Through *in silico* alanisation, we identified a previously uncharacterised residue (D152) in the lid as significant in disrupting the modularity of the 3-way partition. Asp152 is located on 3/10 helix 7 at the boundary between the floppy lid and lid connector helix 8. It has been reported that 3/10 helices are regions of intermediary conformation, suggesting that D152 could play a role in the ‘cracking lid’ dynamics proposed for ADK [34, 35]. Computational evaluation of 17 residue mutations at the D152 site revealed a graded effect due to the different chemical and steric characteristics of the different amino acids (Fig. S3). We confirmed experimentally the relevance of the D152 site by performing single molecule FRET measurements on the wild type and six D152 mutants. Our results showed that the magnitude of the smFRET population shift towards a closed conformation induced by the different mutations is quantitatively correlated with the computed loss of robustness of the large-scale 3-way graph partition.

The largest population shift was observed for D152A, whereas D152E was the least affected and exhibited a similar population equilibrium to the wild type, which is similarly negatively charged. In general, smaller residues with fewer interactions caused a larger shift towards a closed conformation, whilst larger residues with more interactions and/or higher polarity preserved the open conformation. These results indicate that the conformational changes in FRET are a consequence of differing interactions and steric clashes, which are also captured by our atomistic computational analysis. Computationally, the loss of robustness of the large scale communities is traced back to a loss of modular independence of the core and AMP_bd_ domains, which tend to merge. Note that this is a long-range effect: the D152 site is located in the lid, far away from the core-AMP_bd_ interface.

Functionally, all D152 mutations produced full inhibition of the catalytic activity of the enzyme (Fig. S4). Therefore, D152 exemplifies a site where inhibition is not univocally related to the shift towards a closed state. Indeed, D152E has the ability to cycle between the open and closed states, and yet it inhibits catalysis. Rather, our results suggest a widespread reduction in the structural integrity of the domains, which directly affects the chemical geometry of the binding site, and hence an important role for chemical allosteric effects originating from the D152 site. For instance, it is known that D152 forms bonds with a network of arginines (e.g. R150), which have been implicated in catalytic inhibition [28].

Our *in silico* arginine scan revealed a distinct set of mutation sites with large predicted effects on the 3-way partition of ADK. Specifically, we identified V164 in the lid domain, and subsequent smFRET measurements confirmed a strong population shift towards the closed state in V164R. In contrast to D152, however, this shift leads to a proportional reduction of the enzymatic activity, suggesting that the effect of V164R is more directly linked to the minoration of the dynamics of the open:close cycle.

While this manuscript was being finalised, Aviram *et al* published an analysis of smFRET trajectories on *E. coli* ADK measuring domain closure rates as a function of substrate concentration using FRET and a hidden Markov 2-state open-closed model [33]. They propose that numerous cycles of conformational rearrangements, occurring on time scales 100-200 times faster than the turnover rate, are required to find the optimal orientation of substrates. These rearrangements ultimately behave like a ‘bath’ of fluctuations bringing the active site to the right conformation for reaction. Our MS approach probes the structural and atomistic features that could underpin some of these rearrangements, and, through computational mutation and experimental validation, identifies critical residues that can modify the fluctuations and low-lying states in the proposed ‘bath’.

These results highlight the ability of our graph-theoretical approach to identify residues and detailed chemical interactions that influence the protein dynamics at long time and length scales. Noticeably, many of the residues with significant effects under arginine substitution reside in the lid and AMP_bd_ domains, whereas many of the residues significant under alanisation are located in the core subdomain. This difference suggests a role for alanisation in destabilising the hydrophobic core, whereas arginine (due to its charge and size) has a larger effect in the mechanics associated with the lid and binding domains. The remoteness of some of the mutations from the enzyme active site also highlights the impact that single mutations can have by allosterically affecting catalytic function, or being unexpectedly critical for the integrity of functional structure. In particular, the role of Asp152 in the structural integrity of the lid subdomain has so far gone unidentified, yet our analysis shows that it affects both the population conformational equilibrium and its catalytic function. In general, the operation of allosteric mechanisms remains poorly understood [15], and the use of Markov Stability can provide additional information about the intrinsic coupling across scales in protein dynamics affecting their function and structure at long and large scales.

An advantage of scoring mutations using Markov Stability is the retainment of atomistic detail with reduced computational cost, since our framework employs an algorithm that efficiently explores graph partition space to optimise the quality function at high resolution. Large protein complexes, which could take months of MD simulations, can be analysed in minutes. For instance, a full sweep across all Markov times in high resolution for ADK takes 15 minutes on a desktop. Hence full computational mutagenesis scans gauging the effect of point mutations at all scales become computationally accessible.

### Computational methods

#### Protein structures

We analysed two X-ray crystal structures of *Aquifex* ADK (202 residues) obtained by Kern *et al* [24]: (1) the unliganded open conformation (PDB ID: 2RH5) at resolution of 2.48; (2) the closed conformation (PDB ID: 2RGX) at resolution of 1.9, with the atoms of the Ap5A ligand removed.

#### Atomistic network construction

We apply MS to an atomistic protein graph obtained from the three-dimensional protein structure and parameterised with physico-chemical energies (Fig. S1). We summarise briefly the graph construction—for details see [13, 14, 15]. We start from atomistic cartesian coordinates from PDB files. Since X-ray structures do not include hydrogen atoms and NMR structures may be missing some, we use *Reduce* [36] to add any missing hydrogens. The nodes of the graph are atoms and the weighted edges represent interactions, either covalent bonds or weak interactions (hydrophobic, hydrogen bonds or salt bridges) identified using FIRST [37] with a cutoff of 8 for hydrophobic tethers and 0.01 kcal/mol for hydrogen bonds (this value has been optimised in the FIRST software [37]). The edges are weighted by their energies: covalent bonds from standard tables [38]; hydrogen bond energies calculated using the DREIDING force field [39]; hydrophobic bonds calculated using a hydrophobic potential of mean force [40]. The graph is encoded into a weighted adjacency matrix *A*, where element *A*_*ij*_ is the bond energy between atoms *i* and *j* (or zero if there is no interaction). Weighting each edge proportionally to the bond strength encapsulates the idea that an energy fluctuation (as represented by the random walker in the Markov process) is more likely to fluctuate down a chemically stronger bond.

#### Multiscale graph partitioning

Graph partitioning methods aim to partition the nodes of a graph into subgraphs (communities) that are well-connected within themselves and weakly connected to each other. There are multiple ways to define communities, and many methods and criteria to score the resulting partitions [41]. Markov Stability (MS) is a generalised method for identifying communities in graphs at all scales. MS employs a random walk on the graph to define a time-dependent cost function that measures the probability that a random walker is contained within a subgraph over a time scale *t*. If the random walker becomes trapped in particular subgraphs over that particular timescale, this identifies a good partition. As the time scale of the Markov process increases, the method identifies larger subgraphs leading to coarser partitions. Hence MS has the ability to identify intrinsically relevant communities at all scales by using the dynamic scanning provided by the diffusive process, a concept that has been taken advantage of to reveal scale in graphs for purposes of node classification[42] and generalisation of graph centrality [43]. For a detailed description of the method see [2, 17, 18].

Specifically, the random walk is governed by the *N × N* transition matrix *Q* = *D*^−1^*A*, where *N* is the number of nodes in the graph, *A* is the adjacency matrix, and *D* = diag(*A***1**) is the degree matrix where **1** is a vector of ones. *Q* defines the probability of the random walk transitioning from node *i* to node *j*, as given by the discrete-time process:

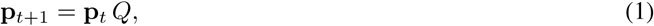

where **p**_*t*_ is a 1 × *N* node vector describing the probability of the random walker to be at each node at time *t*. An associated continuous-time diffusive process in terms of the graph combinatorial Laplacian *L* = *D* − *A* has the time-dependent solution:

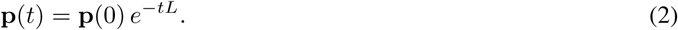

The time *t* is denoted the *Markov time* and is distinct to the biophysical time. The Markov time is a dimensionless quantity related to the diffusive process and acts as a resolution parameter in that it allows for the exploration of the graph at different scales: as the Markov time increases, the partitions become coarser.

A partition of the graph into *c* communities is encoded into a *N* × *c* membership matrix *H*. The goodness of the partition at time *t* under the dynamics governed by *L* is defined in terms of the *c* × *c block auto-covariance* matrix:

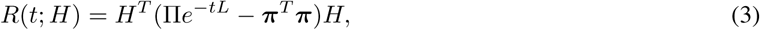

where ***π*** is the stationary solution of (2) and Π = diag(***π***). The element of the matrix [*R*(*t*; *H*)]_*αβ*_ encodes the probability that a random walker starting in community *α* will be at community *β* after time *t*, and the diagonal elements, [*R*(*t, H*)]_*αα*_, indicate the probability of being contained in community *α* over time scale *t*. A good partition *H* maximises the sum of the diagonal elements (i.e., the trace of *R*(*t, H*)); hence we define Markov Stability

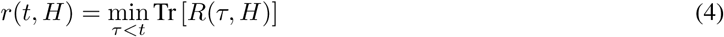

as the cost function to be maximised at every time *t* by searching in the space of partitions *H*:

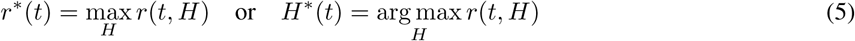

Due to the optimisation in (5) being NP-hard, we use an efficient greedy algorithm known as the *Louvain algorithm* [44], which has been shown to perform well in practice and against benchmarks. The Louvain algorithm is run repeatedly (1000 times for the D152 mutants and 100 times for the full alanine and arginine scans) at each Markov time *t* to obtain an ensemble of optimised partitions {*H*^∗^(*t*)}. We compute the average dissimilarity between partitions in {*H*^∗^(*t*)} using the variation of information [22], denoted VI(*t*). The variation of information is a true metric between clusterings and is closely related to normalised mutual information. Whilst a partition may maximise *r*(*t, H*), it does not necessarily mean that all such partitions are meaningful—it could be the best of a set of bad partitions. Hence we look for robust, highly reproducible partitions. Peaks in VI(*t*) indicate large differences between the partitions found by the optimisation and therefore a lack of robustness; conversely a dip in VI(*t*) indicates consistency in the ensemble of partitions obtained by the Louvain algorithm and therefore a robust partition at that scale *t*. In addition, we search for partitions that are persistent over Markov time (i.e., optimal over extended plateaux) as signalled by low values of VI(*t, t′*), the difference between optimised solutions found at time *t* and *t′*. Partitions that are robust to the optimisation and persistent over Markov time represent relevant intrinsic scales of the graph. Graph partitions at several scales can thus be found in an unsupervised manner without imposing *a priori* the level of resolution.

#### Application to proteins

The inherent complexity of protein dynamics stems from the fact that relevant substructures are dynamically coupled across time scales, from local flexibilities (femtosecond bond vibrations, picosecond to nanosecond methyl rotations, or nanosecond-microsecond loop motions) through to the large collective motions of domains or sub-units moving over time scales spanning up to seconds. Because of its capability to select graph partitions with intrinsic robustness and persistence, as pertains the containment of perturbations within them, MS can be used to identify in an unsupervised manner relevant subgraphs in the protein structure. This process reveals the domain organisation, from small structures such as chemical groups and amino acids (at low Markov times) up to the catalytically important structural domains at long Markov times [13, 14, 23]. While the small scale partitions associated with chemical groups, amino acids, *α*-helices, *β* sheets, etc, are shared by all proteins [13], the partitions found at long Markov times vary across proteins, and can reveal conformational features linked to global structure, dynamics and potentially functional behaviour [45] (Fig. S2).

#### *In silico* mutational analysis

Mutagenesis scans are a widely used experimental approach, in which each residue of a protein is systematically substituted, one at a time, to test its influence on different properties of the protein. Substitution with alanine (i.e., alanisation) eliminates side-chain interactions without introducing steric or electrostatic effects. Mutagenesis scans with other amino acids (e.g., arginine) test the influence of steric and/or electrostatic effects. We have carried out full *in silico* mutagenesis scans of ADK substituting alanine and arginine for each of the 202 ADK residues. Each mutation is introduced directly to the three-dimensional PDB structure using PyRosetta, followed by energy minimisation to produce realistic side-chain packing by running an MD equilibration for 200 ps using GROMACS 5.0 [46]. To evaluate the effect of every mutation [23], we compute the full MS scan of optimal partitions across Markov time and their robustness VI_mut_(*t*) for each of the 202 mutants. We score each mutation by obtaining the averaged difference between the robustness of a relevant partition 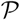 under a mutation *vs.* the wild type

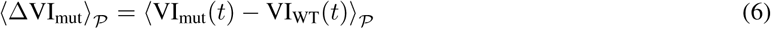

where the average is taken across the Markov time over which the partition 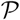 is optimal.

#### Multidimensional Scaling

Multidimensional scaling (MDS) was implemented using the MATLAB mdscale function. The inputs were *D*, an *N × N* dissimilarity matrix (the VI matrix), and the *p*, number of output dimensions (*p* = 2 for a 2-dimensional output). We implemented mdscale using Kruskal’s normalized stress1 criterion. The output *Y* is the euclidean distances between the *N* points which approximate a monotonic transformation of the corresponding dissimilarities in *D*.

### Experimental methods

#### Protein preparation and purification

Protein plasmids of *Aquifex* Adenylate Kinase (ID:18092 Plasmid:peT3aAqAdk/MVGDH) were purchased from AddGene as deposited by ‘Dorothee Kern Lab Plasmids’. The plasmids were already encoded with two cysteine mutations for maleimide conjugation. ADK was expressed in a 1 litre culture BL21 (DE3) cells via inoculation with 1 mM IPTG. BugBuster was used for cell lysis and TCEP and protease inhibitor was added to the lysate. ADK was purified via HIS-tag with a gravi-trap (GE-healthcare), and a PD-10 column was used to remove imidazole and exchange into protein buffer (20 mM TRIS, 5 0mM NaCl). TCEP and protease inhibitor were added throughout the purification process. Alexa 488/Alexa 633-labelled ADK was prepared overnight using 20 *µ*M protein with molar ratio 1:10:10 of protein:Alexa 488:Alexa 633. Excess dye was removed using HIS-tag purification and a PD-10 column. A Typhoon was used to examine the gel of the purified-labelled ADK product and showed no excess fluorophore.

#### Site-specific mutagenesis

Site-specific mutations were introduced into the peT3a-AqAdk/MVGDH plasmid using QuickChange methodology. A PfuUltra High-Fidelity DNA polymerase was used with appropriate oligonucleotides. The plasmids were transformed into XL1-Blue supercompetent cells and a culture was grown for DNA mini-prep. The sequence for each final construct was verified using DNA sequencing (Beckmann Genomics). Successful constructs were transformed into BL21 (DE3) cells for protein growth and extraction. DNA extraction and sequencing was performed again to check the construct in BL21 cells.

#### Single molecule FRET

Freely diffusing single molecules were detected using a home-built, dual-channel confocal fluorescence microscope. Further details on this apparatus can be found in Refs. [47, 48]. An argon ion laser (model 35LAP321, Melles Griot, Carlsbad, CA) with 150 *µ*W at 514.5 nm was used to excite the donor, Alexa 488. An oil-immersion objective (Nikon Plan Apo TIRF 60x, NA 1.45) was employed to collect the donor and acceptor fluorescences which were then separately detected by two photon-counting modules (SPCMAQR14, Perkin-Elmer). Photon detection was recorded by two computer-implemented multichannel scalar cards (MCS-PCI, ORTEC, Canada). Samples were aliquoted for FRET at 50 pM in 200 *µ*l of pH 7.5 FRET buffer (20 mM TRIS, 50 mM NaCl) with 0.3 mg/ml BSA to prevent surface adsorption. A threshold of 35 counts per 500 *µ*s bin for the sum of the donor and acceptor fluorescence signals was used to differentiate single molecule bursts from the background. The background was obtained using independent measurements of buffer solutions without labeled samples and subsequently subtracted from each burst. Background measurements were 0.6 counts and 1.1 counts per 500 *µ*s in the donor and acceptor channels respectively. Apparent FRET efficiencies of each burst were calculated according to: *E*_app_ = *h*_*A*_/(*h*_*A*_ + *h*_*D*_), where *h*_*A*_ and *h*_*D*_ are the acceptor and donor counts, respectively.

To analyse subpopulations within the histogram of the FRET efficiency *E* we fitted a mixture model with Gaussian, *G*(*E*), and four parameter log-normal, *L*(*E*), probability distributions for symmetric and asymmetric peaks, respectively:

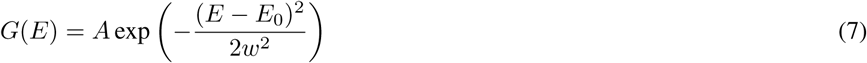

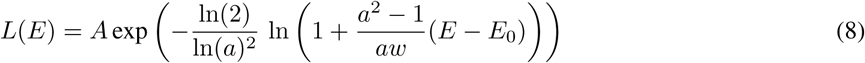

where *A* is the peak amplitude, *E*_0_ is the peak position (mean), *w* is the peak width, and *a* is the asymmetry factor of the peak [49]. The log-normal peak function was used for the donor only population with a fixed width (*w* = 0.15) and asymmetry (*a* = 1.2) and for the high FRET efficiency peaks (*E*_0_ > 0.9). The donor only population (zero peak) was removed after fitting. The remaining FRET-labelled population was fitted with a Gaussian peak function. ADK is known to be a two-state system, hence a two peak distribution (plus the removed log-normal zero peak) is assumed and the histogram was fitted to a mixture of such distributions. The peak positions were identified using the apo, substrate-bound and TMAO osmolyte perturbed structures. The mean and standard deviation was then fixed across all mutant species of the same FRET pair.

#### Kinetic assays

An ADP substrate assay was used to determine the rate of production of ATP. Luminescence via a luciferase reaction with ATP was measured using a time-resolved luminometer with no filter over a 4 hour period. A kinase-glo reagent was used at a total of 100 *µ*l reaction volume in a 96 well-plate. Protein was measured at a 5 nM concentration with 500 *µ*M ADP at constant temperature of 22°C.

## Acknowledgements

This work was funded by an EPSRC Centre for Doctoral Training Studentship (EP/F500416/1) from the Institute of Chemical Biology at Imperial College London awarded to RLP. LY acknowledges funding from the BBSRC (JF20607/2) and the Leverhulme Trust (RPG-2015-345). MB acknowledges funding from the EPSRC award EP/N014529/1 supporting the EPSRC Centre for Mathematics of Precision Healthcare.

## Supplementary Information

**Figure S1:**
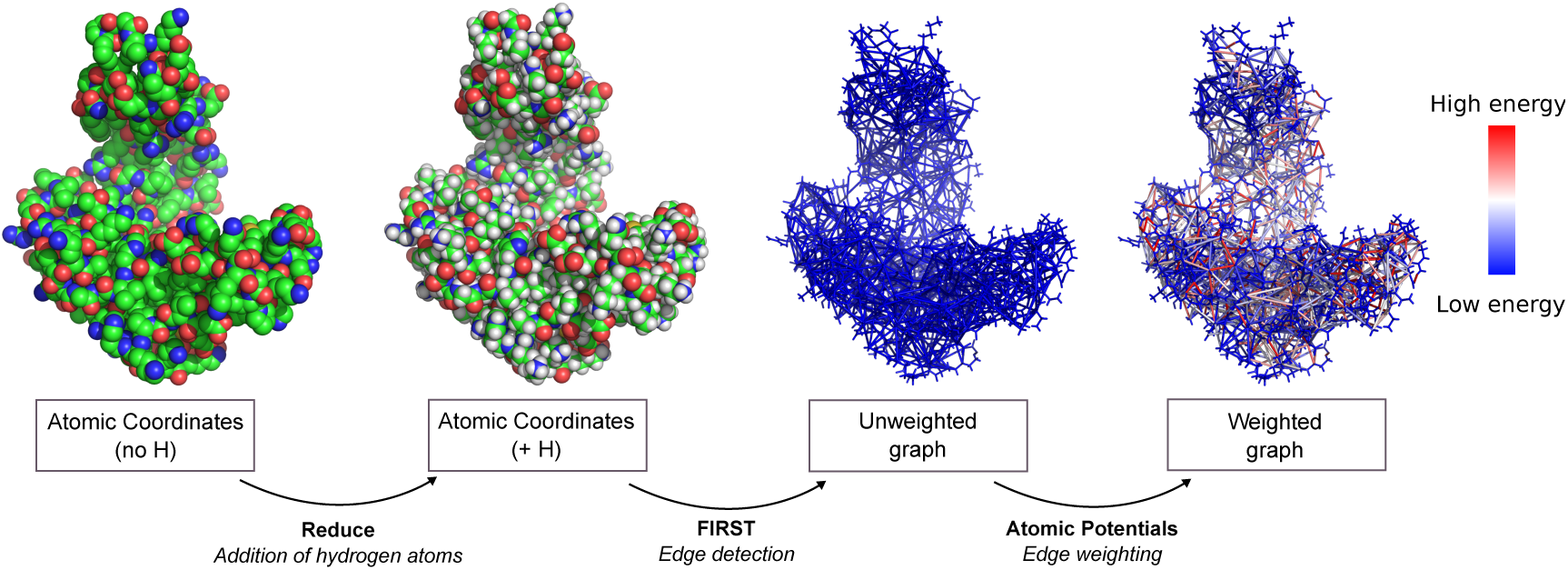
Schematic of the steps of graph construction from a PDB structure. Starting with a PDB file (2RH5 here) the structure is first protonated, then every interaction is identified using FIRST to construct the graph, which is subsequently weighted using appropriate molecular potentials.

**Figure S2:**
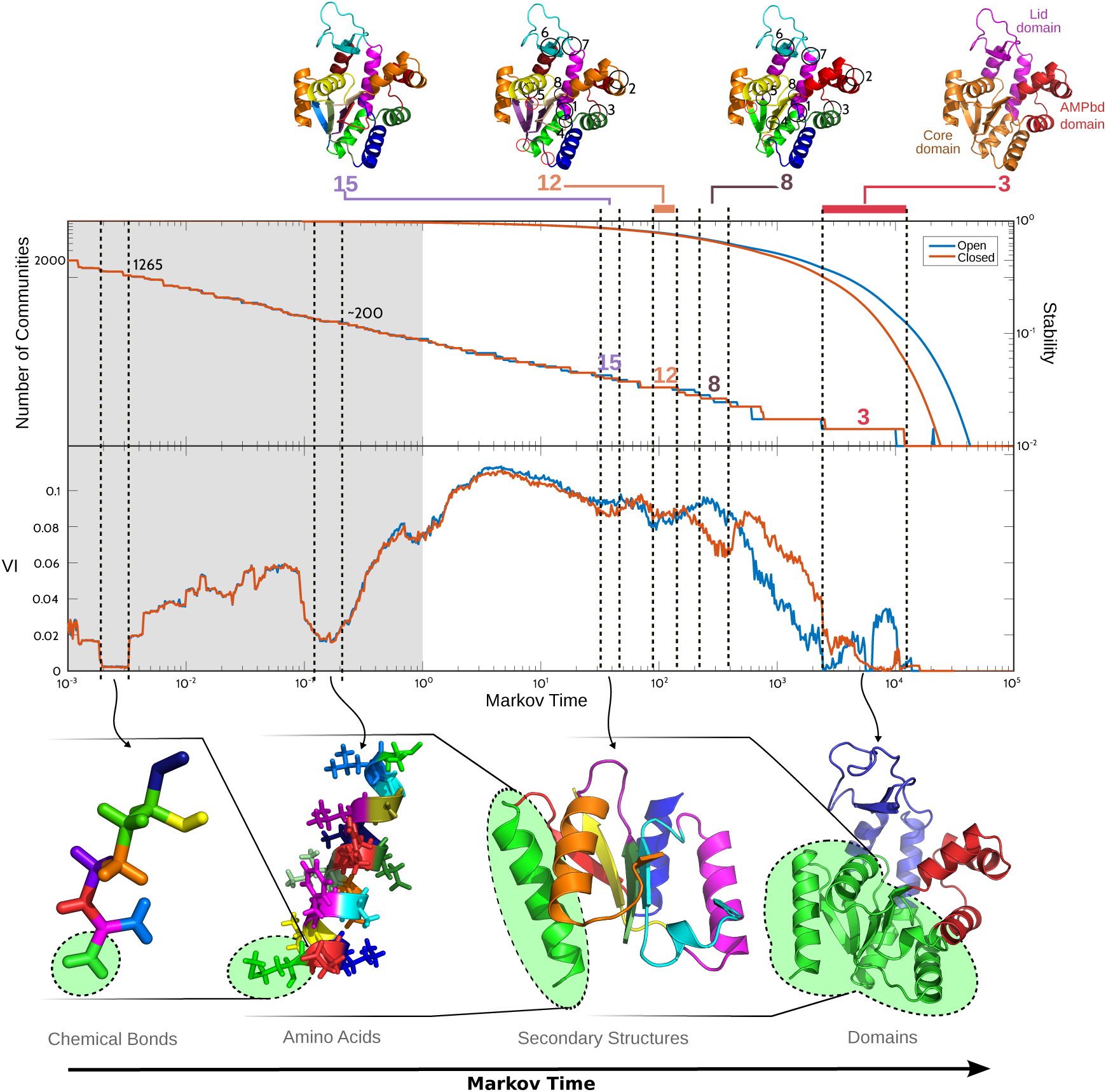
Markov Stability results for the unliganded open conformation (PDB:2RH5, blue) and closed conformation (PDB:2RGX, orange) structures of Aquifex ADK. The plot is an extension of Fig. 1A which only examined Markov time above 10^0^. The unseen region in Fig. 1A is highlighted in grey. We have indicated the presence of the structures found in Fig. 1A. The dips in VI(*t*) found at Markov times 200-300 and 1 × 10^−1^ −2 × 10^−1^ correspond to chemical bonds and amino acids, respectively. Those partitions are common to both structures. The dip at *t* ~ 4 × 10^1^ corresponds to a partition that reveals the secondary structures, such as alpha helices and beta sheets. Beyond 10^2^ the results for the two conformations diverge in both their partitions and robustness VI(*t*), reflecting the differences in their tertiary/quaternary structures. Note that the Markov Stability of the open conformation is higher than that of the closed conformation, highlighting the more robust and modular partitions associated with the open conformation. We find that the VI trajectories look very similar for both proteins until later Markov times where the dynamics of the two states diverges.

**Figure S3:**
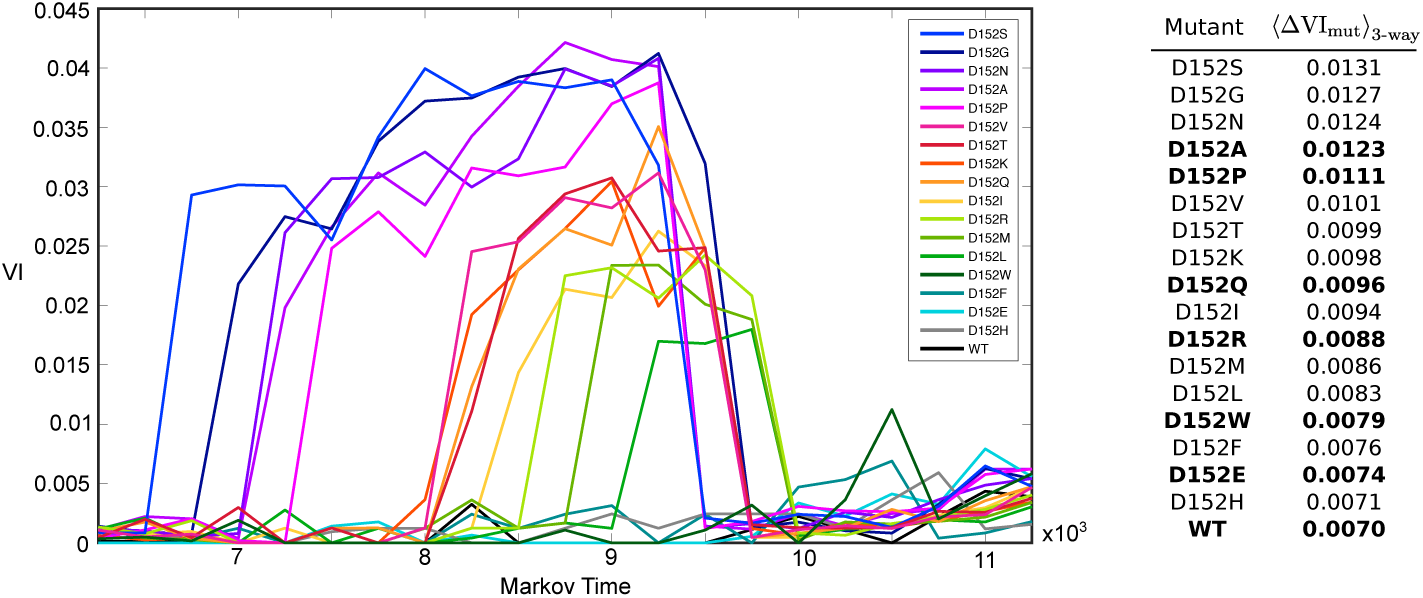
MS analysis of seventeen D152 mutants. The graphs and table are ordered by the effect the mutation on the 3-way partition. Mutants that were experimentally constructed and measured with FRET are shown in bold (see Fig. 3 in the main text).

**Figure S4:**
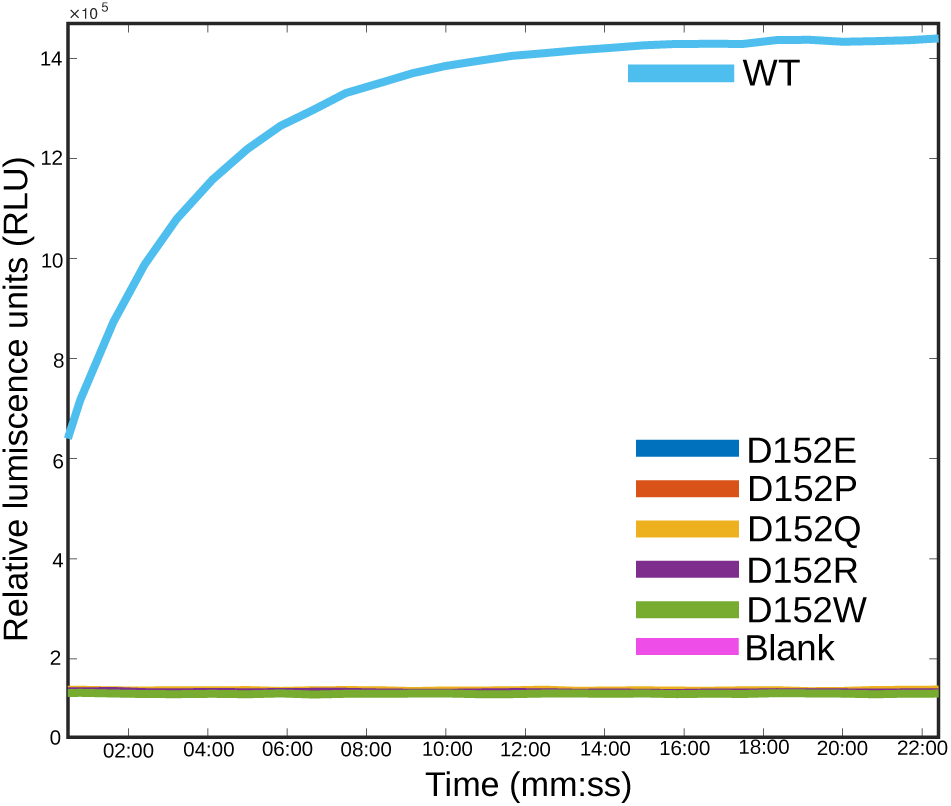
Activity assays of the D152 ADK mutants. A 20 minute window of the 8 hour luminescence activity assay is shown. All D152 mutants exhibit full inhibition relative to the wild type.

**Figure S5:**
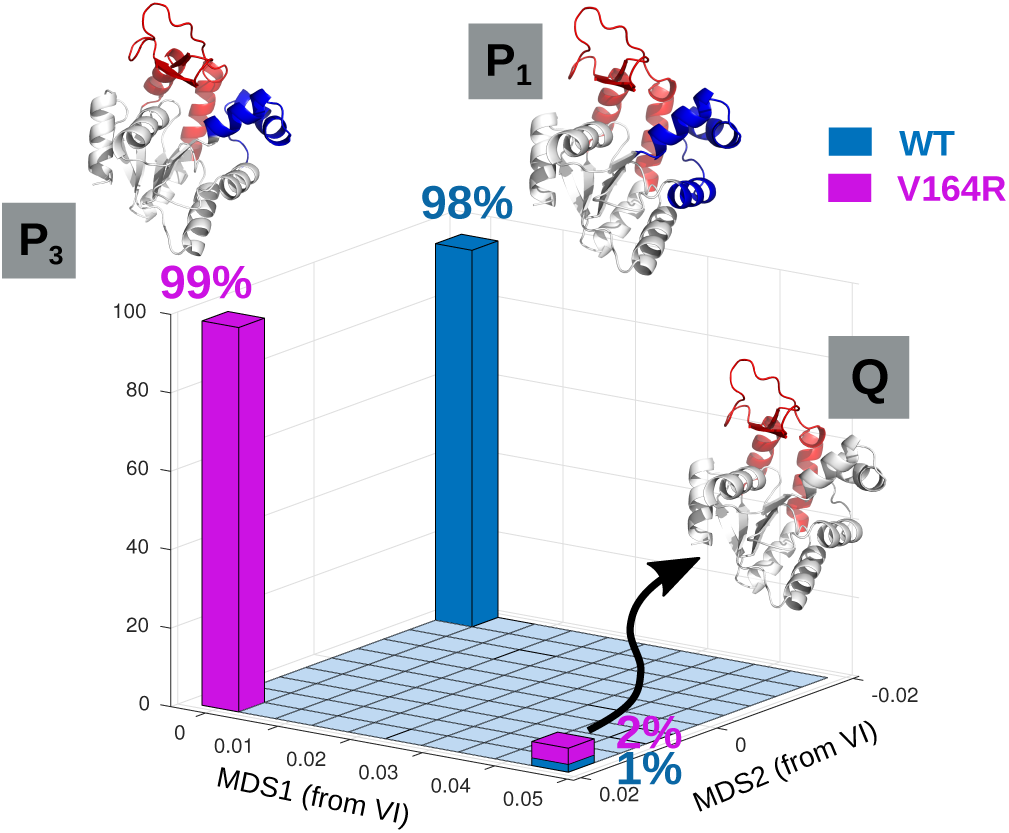
The ensemble of partitions found for V164R shows a loss of modularity of the core and AMP_bd_ domains by shifting alpha helix 4 from AMP_bd_ in the WT to the core in V164R.

**Figure S6:**
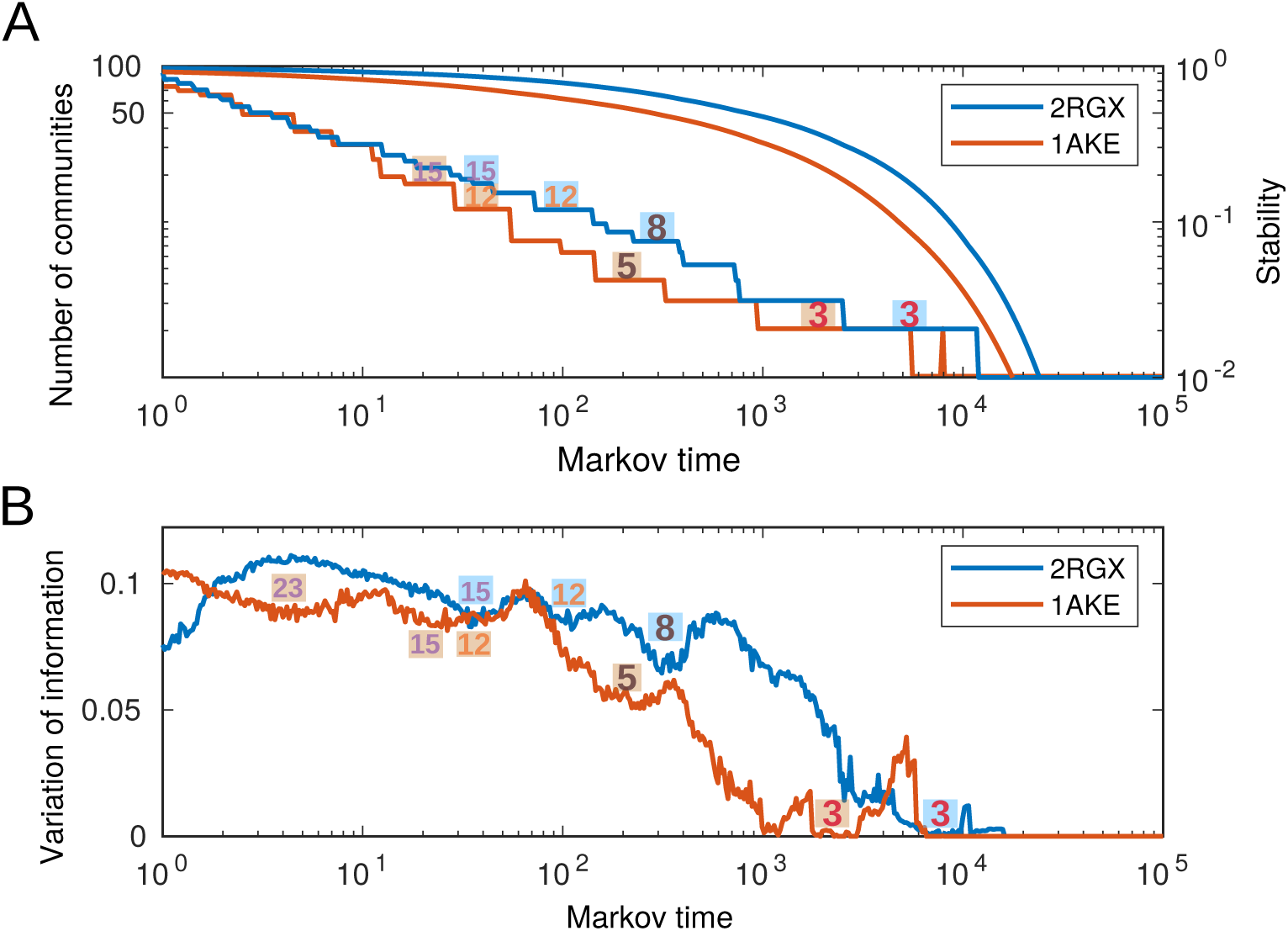
A comparison of the Markov Stability analysis for the E.Coli ADK (PDB ID:1AKE) and Aquifex ADK (PDB ID:2RGX). (a) A comparison of the optimal clustering of the two structures. There are clear differences between the two structures, firstly the E.Coli ADK finds coarse community structures, such as the 3-way partition, much earlier than Aquifex ADK. The global domains of E.Coli ADK appear at earlier Markov times as we would expect with a protein that exhibits faster opening and closing rates [1]. We also identify a 5-way partition as optimal in E.Coli ADK that we don’t observe in Aquifex ADK. Moreover, we don’t observe the 8-way partition in E.Coli ADK that was found in Aquifex ADK. (b) The VI trajectories for the two ADK homologues is used to identify robust partitions. We observe differences in the evolution of *VI* at later Markov times suggest that the underlying structural differences between E.Coli ADK and Aquifex ADK is quite significant.

**Figure S7:**
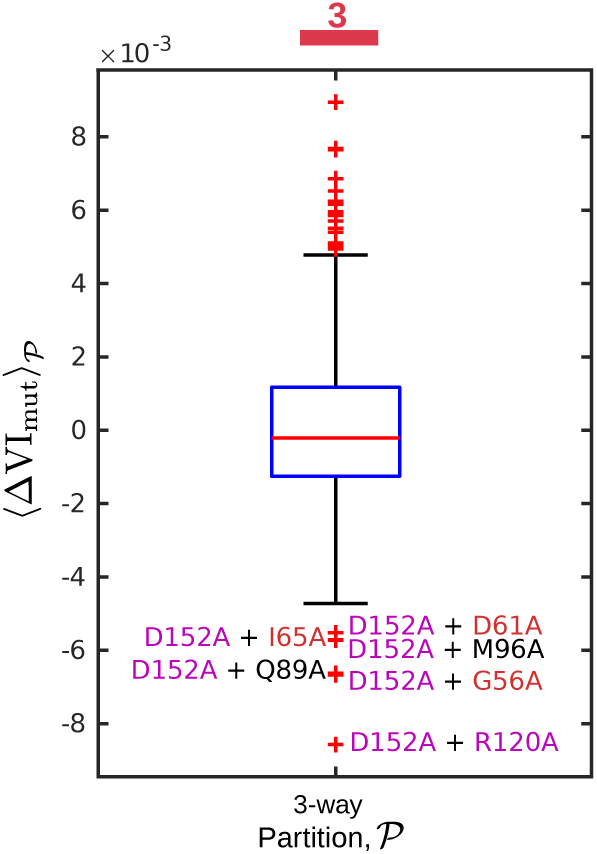
A second alanine scan was peformed on the D152A mutant structure to identify any mutations that would negate the original effect of D152A. The negative values in the boxplot correspond to drops in the VI back towards the original WT. These mutations could be used in further projects to look for paired residues.

**Figure S8:**
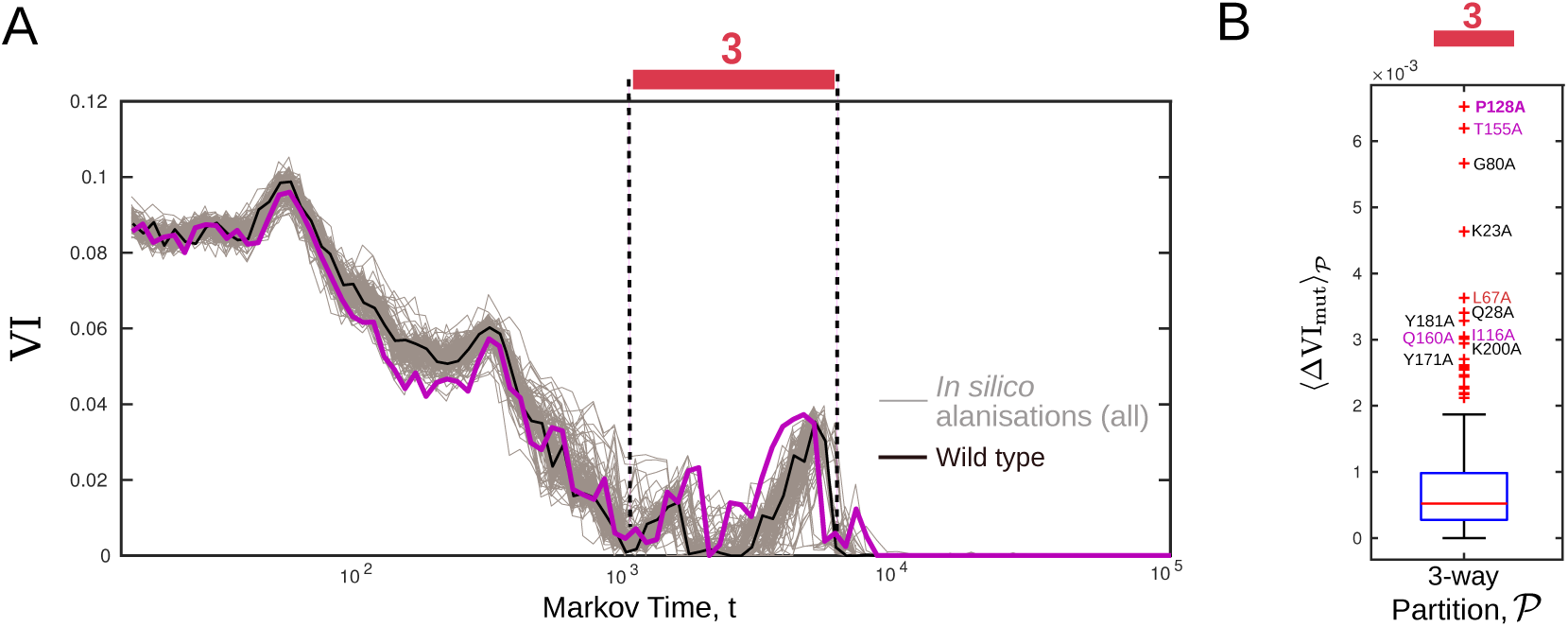
(a) A complete alanine scan and Markov Stability for E.Coli ADK (PDB ID:1AKE) was performed. The *V I* trajectories for the ensemble of mutations are plotted. (b) A boxplot of the mutations that caused the largest fluctuations in *V I* relative to the wildtype. The largest fluctuation during the 3-way partition was observed for P128A which is located in the lid subdomain. In contrast to Aquifex ADK, D152 didn’t appear as important in E.Coli ADK according to this analysis. Mutations in the lid, AMPbd and core subdomains are coloured mauve, red and black respectively.

